# A remodeled RNA polymerase II complex catalyzing viroid RNA-templated transcription

**DOI:** 10.1101/2022.05.31.494195

**Authors:** Shachinthaka D. Dissanayaka Mudiyanselage, Junfei Ma, Tibor Pechan, Olga Pechanova, Bin Liu, Ying Wang

## Abstract

Viroids, a group of plant pathogens, are mysterious subviral agents composed of single-stranded circular noncoding RNAs. Nuclear-replicating viroids exploit host RNA polymerase II (Pol II) activity for transcription from circular RNA genome to minus-strand intermediates, a classic example illustrating the intrinsic RNA-dependent RNA polymerase activity of Pol II. The mechanism for Pol II to accept single-stranded RNAs as templates for transcription remains poorly understood. Here, we reconstituted a robust in vitro transcription system and demonstrated that Pol II also accepts the minus-strand viroid RNA template to generate plus-strand RNAs. Based on this reconstituted system, we purified the Pol II complex on RNA templates for nano-liquid chromatography-tandem mass spectrometry analysis. We identified a remodeled Pol II missing Rpb4, Rpb5, Rpb6, Rpb7, and Rpb9, which contrasts to the canonical 12-subunit Pol II or the 10-subunit Pol II core on DNA templates. This remodeled Pol II is active in transcription with the aid of TFIIIA-7ZF. Interestingly, this remodeled Pol II appears not to require other canonical general transcription factors (such as TFIIA, TFIIB, TFIID, TFIIE, TFIIF, TFIIH, and TFIIS), indicating a different mechanism/machinery regulating RNA-templated transcription. Transcription elongation factors, such as FACT complex, PAF1 complex, and SPT6, were also absent in the transcription complex on RNA templates. We further analyzed the critical zinc finger domains in TFIIIA-7ZF for aiding Pol II activity on RNA template and revealed the first three zinc finger domains pivotal for template binding. Collectively, our data illustrated a distinct organization of Pol II complex on viroid RNA templates, providing new insights into the evolution of transcription machinery, the mechanism of RNA-templated transcription, as well as viroid replication.

## Introduction

Viroids are circular noncoding RNAs that infect crop plants [1, 2]. After five decades of studies, the host machinery for viroid infection has not been fully elucidated [1-3]. There are two viroid families, *Avsunviroidae* and *Popspiviroidae* [4, 5]. Members of *Pospivoirdae* replicate in the nucleus via the rolling circle mechanism (S1 Figure) and rely on the enzymatic activity of RNA polymerase II (Pol II) [1-3]. Specifically, ample evidence support that Pol II activity is critical for the synthesis of oligomer minus-strand or (-) intermediates using viroid circular genomic RNA as templates [1, 6]. However, it remains controversial whether Pol II also uses (-) oligomers as templates for transcription [1, 6].

By and large, RNA polymerases catalyze transcription using DNA templates, which is a fundamental process of life. RNA polymerases are facilitated by a group of general transcription factors to achieve highly regulated transcription, from initiation to elongation and then to termination. Taking Pol II as an example, this 12-subunit complex functions in concert with general transcription factors TFIIA, TFIIB, TFIID, TFIIE, TFIIF, and TFIIH during transcription initiation around DNA promoter regions [7-10]. In general, a minimal system for the promoter-driven transcription requires Pol II and five general transcription factors (TFIIB, TFIID, TFIIE, TFIIF, and TFIIH)[11, 12]. Interestingly, the 10-subunit Pol II core (without Rpb4/Rpb7 heterodimer) is sufficient for transcription elongation [13, 14]. Transcription elongation also requires multiple factors, including TFIIS, TFIIF, DSIF, PAF1 (RNAPII-associated factor 1) complex (PAF1-C), FACT complex (Histone Chaperone), SPT6, etc [15-17].

Since 1974, RNA polymerases have been found to possess intrinsic RNA-dependent RNA polymerase (RdRp) activity to catalyze RNA polymerization using RNA templates [18]. This intrinsic RdRp activity of RNA polymerases was found in bacteria and mammalian cells, as well as exploited by subviral pathogens (i.e., viroids and human hepatitis delta virus (HDV)) for propagation [19-21]. Pol II transcription using viroid or HDV RNA templates can yield RNA products over 1,000 nt in cells, comparable to some products from DNA templates. To ensure such efficient transcription, specific factors are needed. HDV encodes an S-HDAg to promote Pol II activity on its RNA template [22]. Using potato spindle tuber viroid (PSTVd) as a model, we showed that an RNA-specific transcription factor (TFIIIA-7ZF with seven zinc finger domains) interacts with Pol II and specifically enhances Pol II activity on circular genomic RNA template [23, 24]. However, it remains unclear how S-HDAg or TFIIIA-7ZF functions with Pol II for RNA-templated transcription.

Biochemically reconstituted systems have been successfully used to characterize the required factors and functional mechanisms underlying DNA-dependent transcription [25]. However, reconstituted transcription systems using RNA templates often exhibited poor activity. For example, all currently available in vitro transcription (IVT) systems using HDV template have the premature termination issue generating products less than 100 nt [26, 27], which may not reflect the transcription process in cells. We recently established an IVT system for PSTVd [23, 24] that can generate longer-than-unit-length products (more than 360 nt), mimicking the replication process in cells [28].

Using our IVT system, we first confirmed that Pol II and TFIIIA-7ZF function together in transcribing (-) PSTVd oligomers to (+) oligomers. Interestingly, we found that the Pol II complex remaining on (-) PSTVd RNA template has a distinct composition as compared with the 12-subunit Pol II or the 10-subunit Pol II core, via nano-liquid chromatography-tandem mass spectrometry analysis (nLC-MS/MS). Rpb5, Rpb6, and Rpb9 were absent in the remodeled Pol II. Interestingly, Rpb9 is responsible for the fidelity of Pol II transcription. Thus, the absent of Rpb9 may explain the much higher mutation rate of viroid RNA-templated transcription catalyzed by Pol II. Several critical elongation factors for DNA templates, such as the PAF1 complex and SPT6, were also absent in the transcription complex on RNA templates. More importantly, essential general transcription factors (including TFIIA, TFIIB, TFIID, TFIIE, TFIIF, TFIIH, and TFIIS) were all absent in the transcription complex on PSTVd RNA template, clearly demonstrating the distinct regulations between DNA-dependent and RNA-templated transcription. This distinct Pol II retains the catalytic activity to generate (+)-PSTVd oligomers with the aid of TFIIIA-7ZF. We further showed that nearly all seven zinc finger domains of TFIIIA-7ZF are critical for function. In particular, the first three zinc fingers are pivotal for binding with RNA templates. Our findings provide new insights into the organization of Pol II complex on RNA templates, which has profound implications for understanding RNA-templated transcription as well as viroid transcription and its high mutation rates.

## Results

Because PSTVd replication from circular genomic RNA to (-) oligomers and then to plus-strand or (+) oligomers is a continuous process, failure to tease apart each step resulted in controversial data, as evidenced by previous reports [34,41]. To understand whether Pol II can catalyze transcription using (-) viroid oligomers, we established a reconstituted in vitro transcription system using partially purified Pol II from wheat germ [24, 29] and the (-) PSTVd dimer as RNA template. Based on our previous report [24] and the rolling circle replication model (S1 Figure), the linear (+) PSTVd (i.e., the product from IVT assay) cannot serve as a template for further transcription. Therefore, this IVT assay will only focus on the transcription step from (-) PSTVd dimer to (+) PSTVd dimer. As shown in Figure 1a, Pol II has a weak activity in transcribing (-) PSTVd template to full length (+) PSTVd dimer, which is more than 700 nt in length. We then tested the role of the RNA-specific transcription factor (TFIIIA-7ZF) in this transcription reaction by supplying various amounts of the protein. As shown in Figure 1a, TFIIIA-7ZF can significantly increase the RdRp activity (more than 10-fold) of Pol II on the (-) PSTVd dimer template. Therefore, our results indicate that Pol II can accept (-) PSTVd oligomers as a template for transcription, expanding the known natural RNA templates for DdRPs.

**Figure 1.**
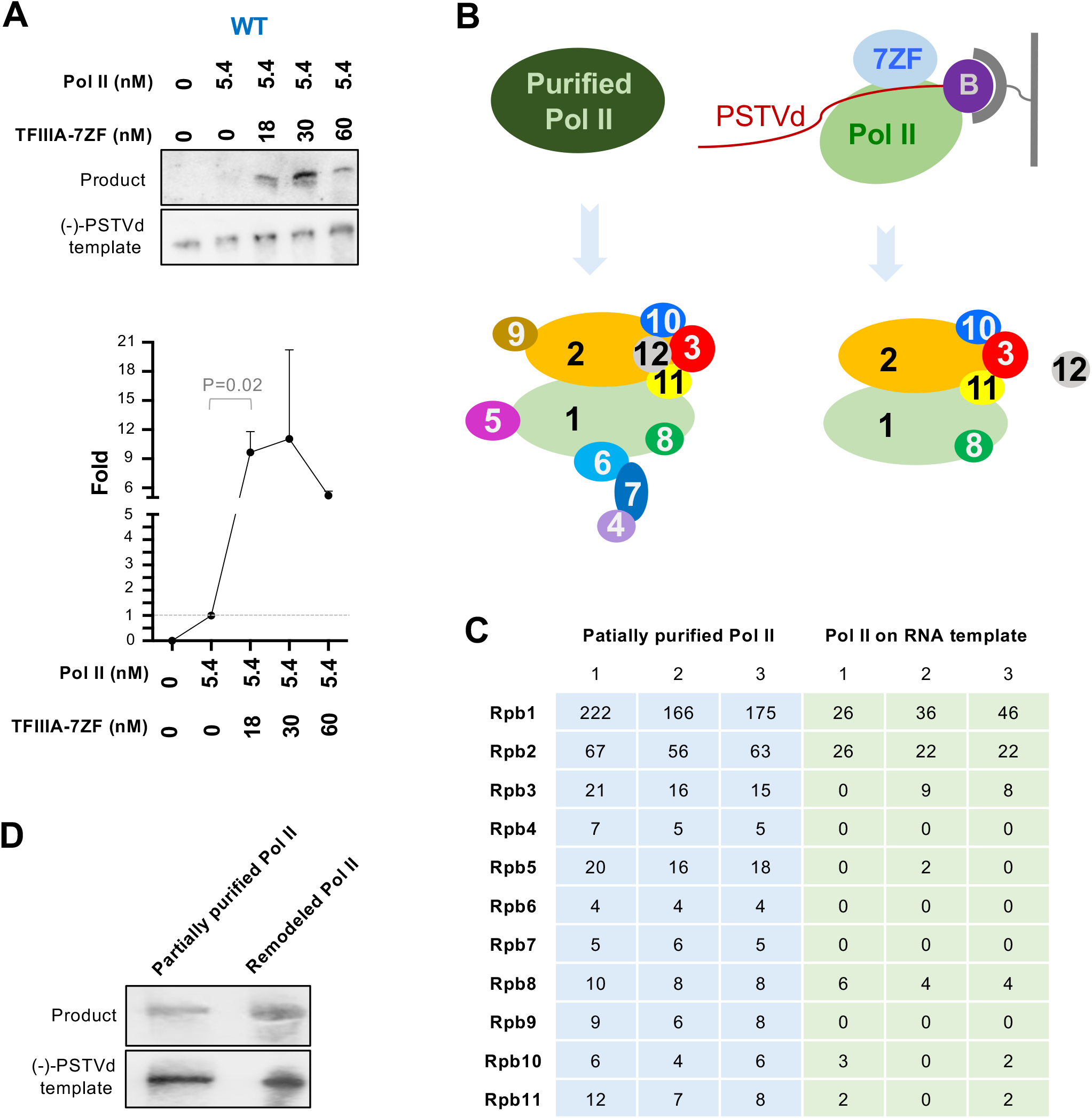
Uncovering the role of TFIIIA-7ZF and Pol II in transcribing (-) PSTVd oligomers. (A) Reconstituted in vitro transcription (IVT) system using partially purified Pol II (from wheat germ), (-) PSTVd dimer RNA (7.8 nM), and various amounts of TFIIIA-7ZF. Sequence-specific riboprobes were used to detect (-) PSTVd templates and (+) PSTVd products (at the position close to dimer PSTVd). Quantification of product intensities was performed using ImageJ. The first lane signal was set as 0 and the second lane signal was set as 1. Data from three replicates were used for graphing the fold increases induced by various amount of TFIIIA-7ZF. (B) Schematic presentation for RNA-based affinity purification followed by nLC-MS/MS identifying protein factors in partially purified Pol II and remodeled Pol II. (C) Peptide counts for each Pol II subunit in all nLC-MS/MS replicates. The summarized nLC-MS/MS data is listed in S1 Table. The original data can be found in S1-S6 datasets. (D) IVT assay demonstrating the activity of remodeled Pol II. The first lane contains free RNA as template, while the second lane contains mixed free RNA and desthiobiothnylated RNA as template. For the second lane, labeled RNA was used first to reconstitute the remodeled Pol II, and the free RNA was then supplied together with NTPs. The reaction condition is described in Methods with details.

We then analyzed the composition of Pol II complex on the viroid RNA template using RNA-based affinity purification. Briefly, a desthiobiotinylated cytidine (Bis)phosphate was ligated to the 3’end of (-) PSTVd dimer, which was then mounted to magnetic streptavidin beads. After sequential supplying of TFIIIA-7ZF and partially purified Pol II, the magnetic beads were washed before elution. Through silver staining, common patterns and distinct bands can be observed between elution fraction and partially purified Pol II (S2 Figure), implying that certain factors may be enriched by or removed from RNA templates. We then performed nLC-MS/MS to reveal the protein factors in partially purified Pol II as well as the Pol II complex remaining on the RNA template (proteins identified in each replicate are listed in S1-S6 Datasets). We identified 11 out of 12 Pol II subunits in partially purified Pol II with high confidence (false discovery rate below 0.01, identified by a minimum of 2 peptides, and 2 peptide-spectrum matches) in all three replicates (Figure 1b and 1c and S1 Table). The smallest subunit Rpb12 (∼6 kDa) was absent, which is possibly caused by sample loss during the size cut-off enrichment of samples for nLC-MS/MS. This issue has been seen in another study [17]. Interestingly, only six subunits were confidently identified in the Pol II complex remaining on the RNA template with high confidence: Rpb1, Rpb2 and Rpb8 were found in all three replicates while Rpb3, Rpb10, and Rpb11 were found in two out of three replicates (Figure 1c and S1 Table). Rpb5 can only be detected in one replicate of Pol II remaining on the RNA template (Figure 1c). Therefore, it is less likely to participate in the Pol II complex on the RNA template. Rpb1 and Rpb2 form the catalytic core of Pol II, while Rpb3, Rpb10, Rpb11 form a subassembly group all critical for Pol II assembly [30]. Since Rpb12 is also a conserved subunit in this subassembly group, we speculate that Rpb12 is also present in the Pol II complex remaining on RNA templates. Rpb8 is an auxiliary factor [30]. Given that the Pol II complex remaining on the RNA template has a distinct composition, we termed it remodeled Pol II hereafter. To test whether the remodeled Pol II has any catalytic activity, we repeated the RNA-based affinity purification followed by IVT. As shown in Figure 1d, this remodeled Pol II indeed possessed transcription activity in generating full-length (+) PSTVd oligomers comparable to partially purified Pol II. It is noteworthy that the product is more than 700 nt in length, comparable to the products generated in cells.

Besides the Pol II complex in the partially purified samples, we also found the presence of several general transcription factors and transcription elongation factors for DNA-dependent transcription. However, they were all absent in the remodeled Pol II in the nLC-MS/MS analysis. For example, we found TFIIF in all three repeats of partially purified Pol II but could not confidently detect it in any of the remodeled Pol II samples (Figure 2 and S1 Table). In addition, TFIIE was found in two out of three repeats of partially purified Pol II but could not be confidently detected in any of the remodeled Pol II samples (S1 Table). Therefore, TFIIE and TFIIF are likely not required for viroid RNA-templated transcription. Noteworthy is that all the rest of the canonical general transcription factors (including TFIIA, TFIIB, TFIID, TFIIH, and TFIIS) were absent in our partially purified or remodeled Pol II. SPT6 is a histone chaperone that interacts with Rpb4/Rpb7 heterodimer during transcription elongation on DNA templates [16]. All Rpb4/Rpb7 and SPT6 were absent in the remodeled Pol II sample (Figure 2 and S1 Table). The FACT complex, including SPT16 and SSRP1-B, is also a histone chaperone assisting Pol II elongation on DNA templates. The FACT complex does not interact with Pol II directly [31], and most of the components also absent in the remodeled Pol II sample (except for the significantly reduced CTR9 in two out of three replicates)(Figure 2 and S1 Table). The PAF1-C, including PAF1, CTR9, LEO1, and RTF1, regulates transcription-coupled histone modifications. PAF1-C components extensively interact with Pol II. PAF1-LEO1 is anchored to the external domains of Rpb2. RTF1 is in proximity to PAF1. The trestle helix in CTR9 binds to Rpb5 and the surrounding region, while the tetratricopeptide repeats interact with Pol II around Rpb11 and Rpb8 [16, 32]. Similarly, the PAF1-C was absent in the remodeled sample (Figure 2 and S1 Table).

**Figure 2.**
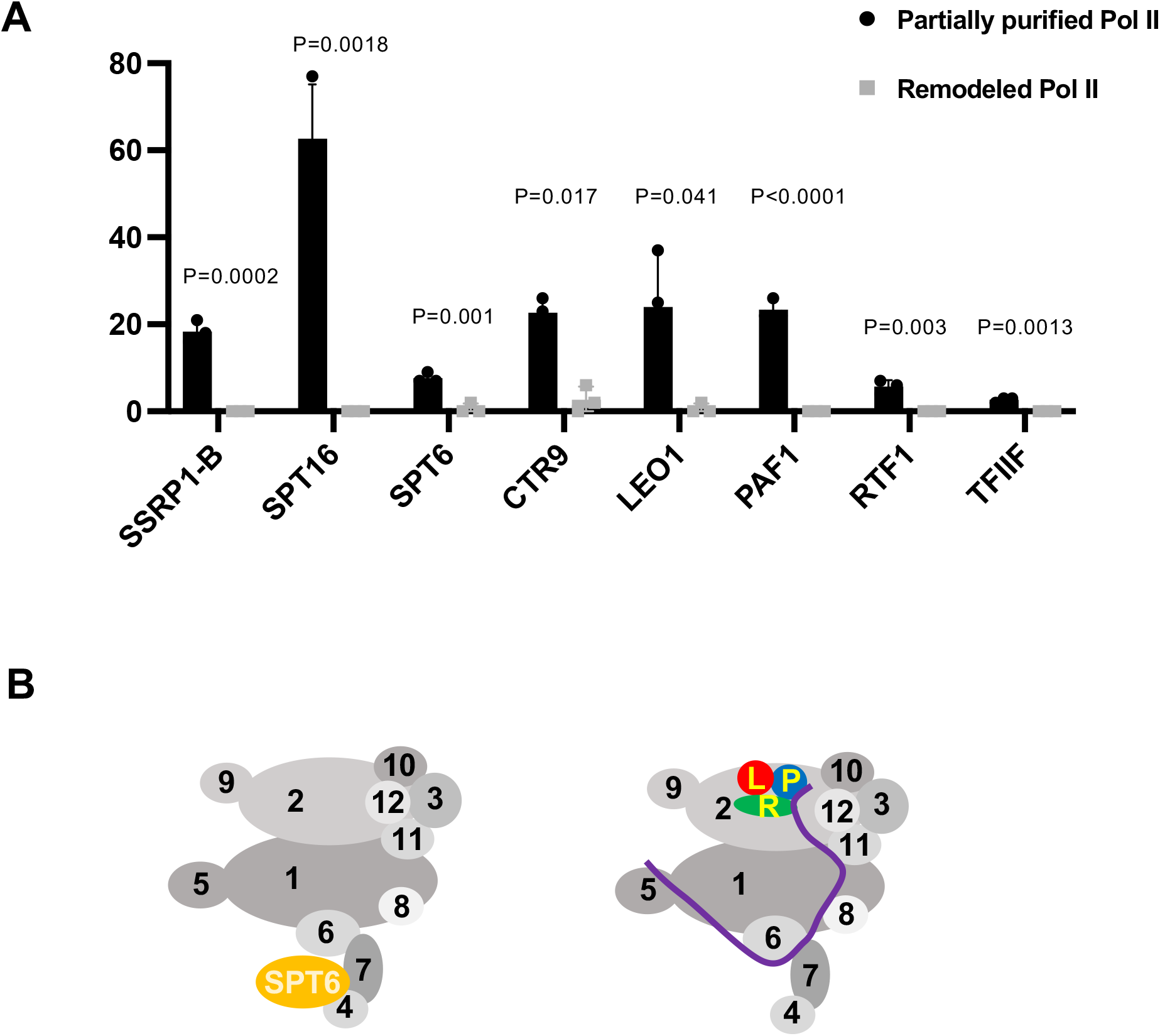
Analyses on transcription elongation factors. (A) Analyses on peptide counts of the FACT complex (SSRP1-B, SPT16), SPT6, PAF1-C, and TFIIF in partially purified Pol II and remodeled Pol II. P values were calculated via two-tailed T-test, a built-in function in Prism. The summarized nLC-MS/MS data are listed in S1 Table. The original data can be found in S1-S6 datasets. (B) Schematic presentation of Pol II interactions with SPT6 and PAF1-C during DNA-dependent transcription, based on [16]. P, PAF1. L, LEO1. R, RTF1. Purple line, CTR9.

Since TFIIIA-7ZF is critical for Pol II to perform transcription using RNA templates, we attempted to identify the functional domain(s) of TFIIIA-7ZF. TFIIIA-7ZF has seven C2H2 type zinc finger domains. We mutated each zinc finger domain by changing the first histidine in the C2H2 domain to asparagine, which is commonly used to disrupt the local structure of a C2H2 motif [33, 34]. We then used those variants for the IVT assay. As shown in Figure 3, all mutants exhibited greatly reduced activity in aiding Pol II transcription on viroid RNA templates. Mutants *zf1, zf2, zf3*, and *zf6* completely lost the activity, while mutants *zf4, zf5*, and *zf7* can increase Pol II activity about two-fold (Figure 3), which is a much weaker activity as compared with more than 10-fold increase stimulated by wildtype (WT) TFIIIA-7ZF (Figure 1A). We then performed the RNA-based affinity purification assay using TFIIIA-7ZF mutants. Since Pol II itself has viroid RNA binding affinity [35], it is not surprising to observe no difference in the amount of the remodeled Pol II on RNA templates with or without the presence of WT TFIIIA-7ZF (Figure 4). Interestingly, *zf1, zf2*, and *zf3* exhibited significantly reduced affinity to (-) PSTVd dimer RNA, which also led to significantly reduced Pol II remaining on RNA templates. The amount of the remodeled Pol II was also reduced, to a lesser extent, in the presence of *zf4, zf5*, or *zf7*, which explains the reduced transcription activity in the corresponding IVT reactions.

**Figure 3.**
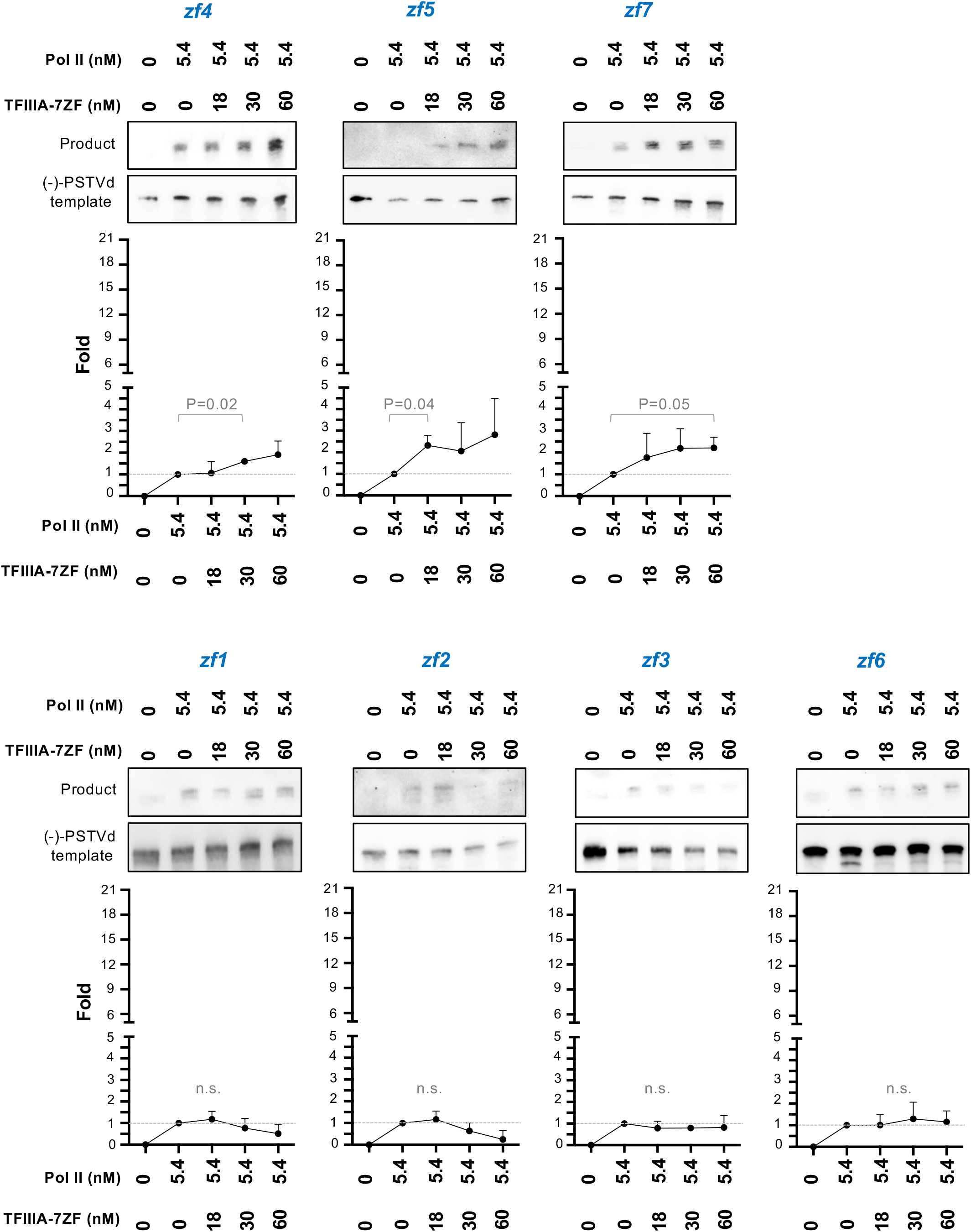
Analyses on the role of TFIIIA-7ZF zinc finger domains in aiding Pol II activity on RNA templates. Reconstituted in vitro transcription (IVT) system using partially purified Pol II, (-) PSTVd dimer RNA (7.8 nM), and various amounts of TFIIIA-7ZF mutants. Sequence-specific riboprobes were used to detect (-) PSTVd templates and (+) PSTVd products (at the position close to dimer PSTVd). Fold changes were analyzed as described in Figure 1. P values for the most significant fold changes as compared with Pol II only samples were listed. n.s., no significant comparisons were identified.

**Figure 4.**
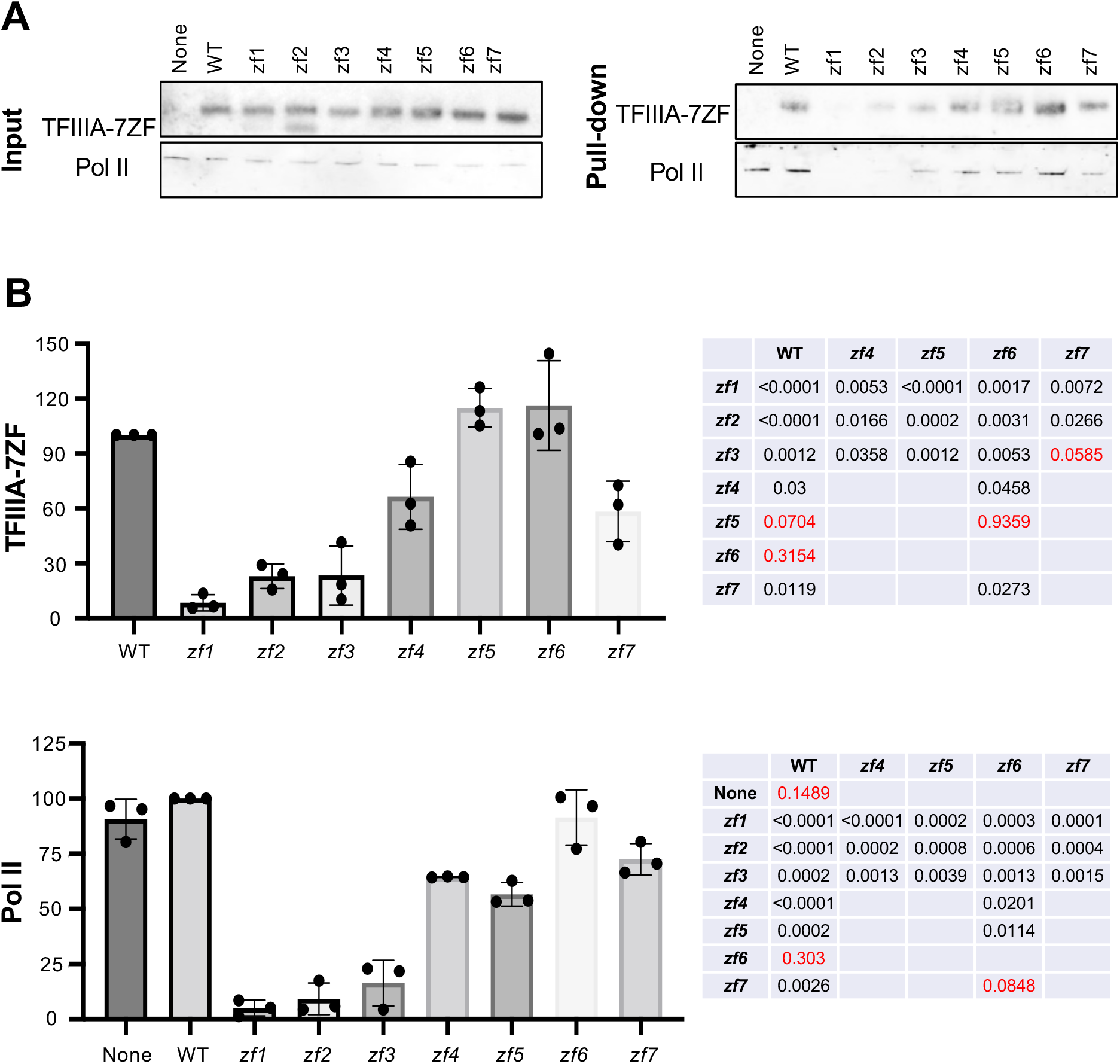
Analyses on the binding ability of TFIIIA-7ZF mutants. WT and mutants TFIIIA-7ZF proteins were used for RNA-based affinity purification in the presence of partially purified Pol II. (A) Immunoblots for input and RNA-bound TFIIIA-7ZF (anti-TFIIIA) and the Rpb1 subunit of Pol II (8WG16). None, no TFIIIA-7ZF protein supplied. (B) Quantification of immunoblotting results in (A). Protein signals in Pull-down blots were normalized to the corresponding signals in the Input blots. The normalized signals in WT (TFIIIA and Rpb1) were set as 100. Three replicates were performed to quantification and statistical analyses. Two tailed T-test was used to calculate P values (listed in tables), by using the built-in function in Prism.

## Discussions

Using a robust IVT platform, we found that a remodeled Pol II and TFIIIA-7ZF can efficiently utilize (-) PSTVd dimer for RNA-templated transcription. TFIIIA-7ZF significantly enhances Pol II transcription activity on RNA templates. This remodeled Pol II represents a new organization of functional polymerase complex. In this remodeled Pol II, we observed the catalytic core (Rpb1 and Rpb2), a subassembly group (Rpb3, Rpb10, Rpb11, likely Rpb12 as well), and an assembly factor Rpb8. Rpb4/Rpb7 heterodimer, Rpb6, Rpb9, and likely Rpb5 were absent in the remodeled Pol II. Rpb4/Rpb7 heterodimer is not essential for elongation and is not included in the Pol II core [13, 14]. Rpb6 is involved in contact with TFIIS and TFIIH [36, 37], neither of which were present in the transcription complex on our viroid RNA template. Furthermore, we showed that TFIIS is not required for PSTVd replication [24]. Rpb9 is critical for Pol II fidelity by delaying NTP sequestration [38, 39]. Pol II fidelity is also regulated by TFIIS [40, 41]. Interesting, both Rpb9 and TFIIS were absent in the remodeled Pol II samples, which is in line with the observation that nuclear replicating viroids have a much higher mutation rate than cellular Pol II transcripts [42]. Rpb5 is proposed to make contacts with DNA promoters and coordinate the opening/closing of the Pol II DNA cleft [43-47], which is not involved in RNA-templated transcription.

It has been proposed that an RNA polymerase may have evolved to transcribe RNA templates first and then transitioned to use DNA templates in modern life forms [30]. Interestingly, Rpb4/Rpb7 heterodimer is present in all archaeal and eukaryotic cells but not in bacteria [10, 48], further suggesting that organization changes occurred in RNA polymerases during the course of evolution. The remodeled Pol II identified here only retains a reduced set of factors, while most of the missing subunits are absent in bacterial RNA polymerases (i.e., Rpb4, Rpb5, Rpb7, and Rpb9). Our discovery of a remodeled Pol II actively transcribing RNA templates may provide a handle to further explore the functional evolution of transcription machinery.

The high-resolution crystallographic structure of Pol II core-Rpb4/Rpb7-TFIIS showed that Pol II utilizes the same active site for interacting DNA and RNA templates, revealed by using chimeric RNA templates [49]. Later, one study using a chimeric RNA template containing a HDV fragment sequence suggested that multiple general transcription factors (TFIIA, TFIIB, TFIID, TFIIE, TFIIF, TFIIH, and TFIIS) may be potentially involved in the initiation of RNA-templated transcription [50]. However, neither experimental system could yield long RNA products, suggesting that those transcription complexes might not be optimal for subviral RNA templates. In the transcription complex on PSTVd RNA template, we could not detect the presence of TFIIE or TFIIF, despite that they were both identified confidently in at least two out of three replicates in partially purified Pol II (S1 Table). It is noteworthy that other Pol II-associated general transcription factors (TFIIA, TFIIB, TFIID, TFIIH, and TFIIS) were also absent in partially purified Pol II and remodeled Pol II samples. Although we cannot rule out the possibility that minute amount of general transcription factors were remaining in our samples below the detection capacity of nLC-MS/MS, it is unlikely for them to play a role in a stoichiometric ratio to Pol II resembling the transcription complex on DNA templates. Therefore, our observation argues that those general transcription factors for DNA-dependent transcription are not required in the transcription complex on RNA templates, at least for viroid RNA templates. Hence, the results outlined a distinct organization of transcription complex on RNA templates.

During transcription elongation on DNA templates, multiple auxiliary factors play regulatory roles in cells. For example, the FACT complex and PAF1 complex regulate transcription elongation. However, those factors may not be needed for in vitro transcription [13, 14]. In line with the previous observations, PAF1 complex and most of the FACT components (except for significantly reduced CTR9) were absent in the transcription complex on PSTVd RNA templates (Figure 2 and S1 Table). Therefore, it awaits future investigations to understand whether those factors are involved in RNA-templated transcription.

All seven zinc finger domains of TFIIIA-7ZF are critical for the function in aiding RNA-templated transcription. The first three zinc finger domains are pivotal for RNA template binding. Interestingly, Pol II exhibited weaker affinity to RNA templates in the presence of either *zf1, zf2*, or *zf3* mutant. It is intuitive to speculate that free *zf1, zf2*, and *zf3* sequestered Pol II and prevented Pol II binding to RNA templates, leading to greatly reduced transcription activity. It is unclear why the *zf6* mutant also greatly diminished Pol II activity in generating full-length product as the amount of *zf6* and Pol II remaining on the RNA template resembles that in the reaction with WT TFIIIA-7ZF. Since our system only tested TFIIIA-7ZF and Pol II binding to the RNA templated before reaction initiates, reasonable speculation is that *zf6* might be critical for transcription initiation or even elongation.

Our reconstituted IVT system is robust for exploring the factors and functional mechanism underlying viroid RNA-templated transcription, particularly for studying transcription initiation and elongation. However, the current system does not have a regulated termination process as the transcription is terminated by template run-off. A modified system is needed in the future to serve the purpose of studying transcription termination on RNA templates. In addition, structural analyses of the remodeled Pol II/TFIIIA-7ZF complex on viroid RNA templates may provide more insights into the regulation and mechanism underlying RNA-templated transcription.

## Materials and Methods

### Molecular constructs

We have previously reported WT TFIIIA-7ZF cloned from *Nicotiana benthamiana* in bacteria expression vector pTXB1 (New England Biolabs, Ipswich, MA) [23]. The TFIIIA-7ZF mutants were generated via site-directed mutagenesis using the WT TFIIIA-7ZF in pTXB1 as the template (See S2 Table for primer information). We have also reported the pInt95-94(-) and pInt95-94(+) constructs for generating PSTVd probes to detect sense and antisense PSTVd RNAs, respectively [23]. PSTVd dimer construct was reported previously [51]. All constructs have been verified by Sanger sequencing.

### Protein purification

The protocol for recombinant protein purification was based on our reported protocol [23]. Various recombinant TFIIIA-7ZF proteins with an intein-chitin binding domain (CBD) tag were overexpressed using the *Escherichia coli* BL21(DE3) Rosetta strain (EMD Millipore, Burlington, MA). For each construct, about 500 ml bacterial culture was collected and re-suspended. After sonication with Bioruptor (Diagenode, Denville, NJ), samples were subjected to centrifugation at 15,000X *g* for 1 h at 4°C. The cell lysate was collected and incubated for 1 h with 2 ml of 50% slurry of chitin resin (New England Biolabs) before loading onto an empty EconoPac gravity-flow column (Bio-Rad Laboratories, Hercules, CA). After washing, resin was incubated for 18 h at 4°C in a cleavage buffer [20 mM Tris-HCl (pH 8.5), 500 mM NaCl, 50 mM dithiothreitol, and 5 µM ZnSO_4_]. Fractions containing tagless proteins were dialyzed against 20 mM HEPES, pH 7.5, 200 mM NaCl, 50 µM ZnSO_4,_ and 5 mM DTT. Protein concentrations were estimated by Coomassie Brilliant Blue staining of an SDS-PAGE gel using as reference standards bovine serum albumin of known concentrations.

Purification of Pol II from wheat germ was carried out following our published protocol [24]. All operations were performed at 4°C, and all centrifugations were carried out for 15 min. One hundred grams raw wheat germ (Bob’s Red Mill, Milwaukie, OR) was ground in a Waring Blender with 400 mL of buffer A [50 mM Tris-HCl pH 7.9, 0.1 mM EDTA, 1 mM DTT, and 75 mM (NH_4_)_2_SO_4_]. The resulting homogenate was diluted with 100 mL of buffer A and followed by centrifugation at 15,000× *g*. Supernatant was filtered through one layer of Miracloth (MilliporeSigma, Burlington, MA). The resulting crude extract containing Pol II was precipitated by an addition of 0.075 volume of 10% (v/v) Polymin P with rapid stirring. The resulting mixture was subject to centrifugation at 10,000X *g*. The pellet was washed with 200 mL of buffer A. The insoluble fraction, which contains Pol II, was resuspended with the buffer B [50 mM Tris-HCl pH 7.9, 0.1 mM EDTA, 1 mM DTT, and 0.2 M (NH_4_)_2_SO_4_]. The resulting suspension was centrifuged at 10,000X *g* to remove insoluble pellets. (NH_4_)_2_SO_4_ precipitation was carried out by slowly adding 20 g of solid (NH_4_)_2_SO_4_ per 100 mL of the above supernatant with stirring. Mixture was centrifuged, and the pellet was dissolved in buffer C (0.05 M Tris-HCl pH 7.9, 0.1 mM EDTA, 1 mM DTT, 25% ethylene glycol) plus 0.1% Brij 35 (Thermo Fisher Scientific, Waltham, MA) to make final (NH_4_)_2_SO_4_ concentration 0.15 M. The (NH_4_)_2_SO_4_ concentration was determined by conductivity. The resulting solution was applied to DEAE Sepharose FF (GE Healthcare Life Sciences, Pittsburgh, PA) equilibrated with buffer C plus 0.15 M (NH_4_)_2_SO_4_. Then column was washed with five bed volume with buffer C containing 0.15 M (NH_4_)_2_SO_4_. Finally, bound Pol II was eluted with buffer C containing 0.25 M (NH_4_)_2_SO_4_. Fractions containing Pol II were pooled. The (NH_4_)_2_SO_4_ concentration was adjusted to 75 mM by conductivity. Resulting solution was applied to SP Sepharose FF (GE Healthcare Life Sciences) equilibrated with the buffer C containing 75 mM (NH_4_)_2_SO_4_. After washing the column with the same buffer, Pol II was eluted using the buffer C containing 0.15 M (NH_4_)_2_SO_4_. Eluted fractions containing Pol II were pooled. Ethylene glycol (VWR Chemicals BDH, Radnor, PA) was added to a final concentration of 50% (v/v) before storing at −20°C.

### In vitro transcription assay

Pol II-catalyzed in vitro transcription was carried out based on our recently developed protocol [23, 24]. BSA (New England Biolabs) and TFIIIA-7ZF were treated with 1 unit of Turbo DNase (Thermo Fisher Scientific) for 10 min at 37°C. Then, 0.39 pmol (-) PSTVd dimer, 0.27 pmol partially purified Pol II, pretreated BSA (4 μM final concentration) and various amounts of TFIIIA-7ZF were incubated at 28°C for 15 min. The reaction system was adjusted to contain 50 mM HEPES-KOH pH 7.9, 1 mM MnCl_2_, 6 mM MgCl_2_, 40 mM (NH_4_)_2_SO_4_, 10% (v/v) glycerol, 1 unit/μL SuperaseIn RNase inhibitor (Thermo Fisher Scientific), 0.5 mM rATP, 0.5 mM rCTP, 0.5 mM rGTP, 0.5 mM rUTP. Transcription reactions (50 μL) were incubated at 28°C for 4 h. About 0.8 U/μL proteinase K (New England Biolabs) was applied to terminate the reaction by incubation at 37°C for 15 min, followed by incubation at 95°C for 5 min. The Pol II-catalyzed in vitro transcription assay was repeated three times for each TFIIIA-7ZF variant. For the IVT assay in Figure 1D, TFIIIA-7ZF and partially purified Pol II were subsequentially bound to immobilized desthiobiotinylated RNA templates (see the section below for details). After washing twice, additional RNA templates without desthiobiotinylation were supplied together with NTPs. The reaction was then performed the same as abovementioned. This assay was repeated twice.

### RNA-based affinity purification

Using Pierce RNA 3’ end desthiobiotinylation kit (Thermo Fisher Scientific, Waltham, MA, USA), 50 pmol of PSTVd dimer RNA was labeled following the manufacturer’s instructions. Labeled RNA was purified using MEGAclear kit (Thermo Fisher Scientific, Waltham, MA, USA) and heated at 65°C for 10 min followed by incubation at room temperature for 12 min. Labeled RNA was bound to the 50 μL of streptavidin magnetic beads (Thermo Fisher Scientific, Waltham, MA, USA). Magnetic beads were collected and washed twice with equal volume of 20 mM Tris-HCl, pH 7.5. Beads were subsequently washed with reaction buffer containing, 50 mM HEPES-KOH pH 8, 2 mM MnCl_2_, 6 mM MgCl_2_, 40 mM (NH_4_)_2_SO_4_, 10% glycerol. DNase treated 150 pmol of recombinant TFIIIA-7ZF was incubated with RNA-bound beads in a 50 μL reaction at 28°C for 15 min. Then, 27 pmol of partially purified Pol II was added to the reaction and incubated at 28°C for another 15 min. Next, beads were washed twice with wash buffer (20mM Tris-HCl, pH 7.5, 10mM NaCl, 0.1% Tween-20) and bound proteins were eluted with 1X SDS-loading buffer by heating 95°C for 5 min.

### RNA gel blots

Detailed protocol has been reported previously [52]. Briefly, after electrophoresis in 5% (w/v) polyacrylamide/8 M urea gel for 1 h at 200 V, RNAs were then transferred to Hybond-XL nylon membranes (Amersham Biosciences, Little Chalfont, United Kingdom) by a Bio-Rad semi-dry transfer cassette and were immobilized by a UV-crosslinker (UVP, Upland, CA). The RNAs were then detected by DIG-labeled UTP probes. PSTVd RNAs were prepared as described before [23]. *Sma*I-linearized pInt95-94(-) and pInt95-94(+) were used as templates for generating probes, using the MAXIclear kit (Thermo Fisher Scientific). The DIG-labeled probes were used for detecting PSTVd RNAs.

### Immunoblots

Protein samples were separated on an SDS-PAGE gel, followed by transferring to nitrocellulose membrane (GE Healthcare Lifesciences) using the Mini-PROTEAN Tetra Cell (Bio-Rad Laboratories). After 1 h incubation with 1% (w/v) nonfat milk in 1X TBS (50 mM Tris-HCl, pH 7.5, 150 mM NaCl) at room temperature, membranes were incubated with primary antibodies overnight at 4°C. After three washes with 1X TBST (50 mM Tris-HCl, pH 7.5, 150 mM NaCl, 0.1% Tween 20), HRP-conjugated secondary antibodies were added. Membrane was then washed three times with 1X TBST and incubated with HRP substrates (Li-COR Biosciences, Lincoln, NE). The signals were detected with ChemiDoc (Bio-Rad Laboratories).

For immunoblotting, polyclonal antibodies against TFIIIA were diluted as 1:2,000 and the monoclonal 8WG16 antibodies (Thermo Fisher Scientific) were diluted at 1:1,000. HRP-conjugated anti-mouse serum (Bio-Rad) was diluted at 1:5,000. HRP-conjugated anti-rabbit serum (Thermo Fisher Scientific) was diluted at 1:3,000. For silver staining, we followed instructions of Silver BULLit kit (Amresco, Solon, OH).

### Nano-liquid chromatography-tandem mass spectrometry analysis (nLC-MS/MS)

Prior mass spectrometry, samples were subjected to in-solution digestion. Briefly, reduction treatment (100 mM dithiothreitol and 15 min incubation at 65°C) and alkylation treatment (100 mM iodoacetamide / 45 min incubation at room temperature) were followed by 16 hr incubation at 37°C with sequencing grade trypsin (Promega, Madison WI). Tryptic peptides were acidified with formic acid, lyophilized and stored at −80°C. As described previously [53], two micrograms of digested protein were subjected to nLC-MS/MS analysis using the LTQ-Orbitrap Velos mass spectrometer (Thermo Fisher Scientific) directly linked to the Ultimate 3000 UPLC system (Thermo Fisher Scientific), with following modification: mass spectra of intact and fragmented peptides were collected in the orbitrap and linear ion trap detector, respectively. All data files were deposited to PRIDE database [54] (accession PXD033736). The .raw mass spectral files were searched using the SEQUEST algorithm of the Proteome Discoverer (PD) software version 2.1 (Thermo Fisher Scientific). Tolerances were set to 10 ppm and 0.8 Da to match precursor and fragment monoisotopic masses, respectively. The Triticum NCBI Ref protein database (as of February 2022, with 122,221 entries) and its reversed copy served as the target and decoy database, respectively, to allow calculation of False Discovery Rate (FDR). All proteins presented in results were filtered by FDR<1%, and identified by minimum of 2 peptides and 2 PSMs (peptide-spectrum matches) in each replicate. The PD result data files showing peptide/protein-ID relevant parameters for each individual replicate are given in Supplementary Material.

## Supporting information

S2 FigureFigure

## Acknowledgements

This work was supported by US National Science Foundation (MCB-1906060 and MCB-2145967 to YW), US National Institute of General Medical Sciences (1R15GM135893 to YW), and NIH MS-IDeA Network of Biomedical Research Excellence award 5P20GM103476-19. The mass spectrometry proteomics analysis was performed at the Institute for Genomics, Biocomputing and Biotechnology, Mississippi State University, with partial support from Mississippi Agricultural and Forestry Experiment Station. We are grateful for the constructive comments from Donna Gordon at Mississippi State University.

## Accession Numbers

The mass spectrometry proteomics data have been deposited to the ProteomeXchange Consortium via the PRIDE partner repository with the dataset identifier PXD033736.

Please note, the data will be publicly available upon acceptance of the manuscript. Reviewers can access the data by using the following login information: Username; Password: yThN9ExZ

## Author Contributions

Y.W. conceived and designed the experiments. S.D.DM., T.P., B.L., and Y.W. wrote the manuscript. S.D.DM., T.P., and O.P. performed experiments. S.D.DM., J.M., and Y.W. prepared key materials. S.D.DM., T.P., B.L., and Y.W. analyzed the data.

