## Supplementary figures and images for "A remodeled RNA polymerase II complex catalyzing viroid RNA-templated transcription"

### S1 Figure

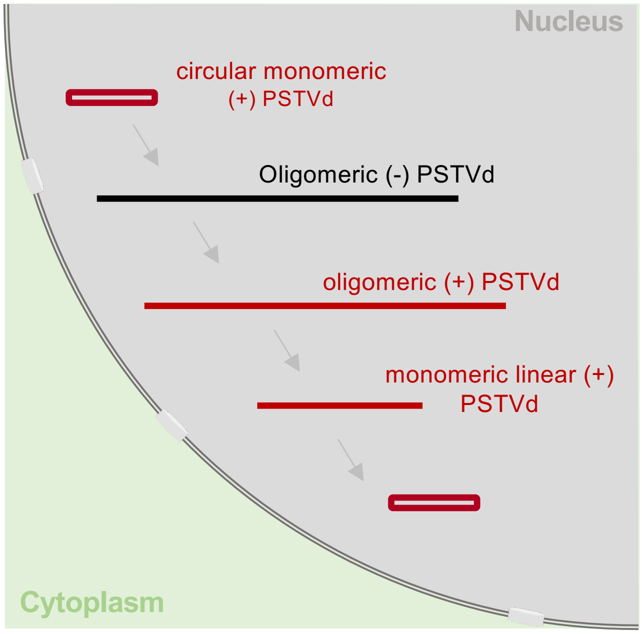


**S1 Figure.** Rolling circle replication of PSTVd.
