## Supplementary material for "A remodeled RNA polymerase II complex catalyzing viroid RNA-templated transcription": S2 FigureFigure

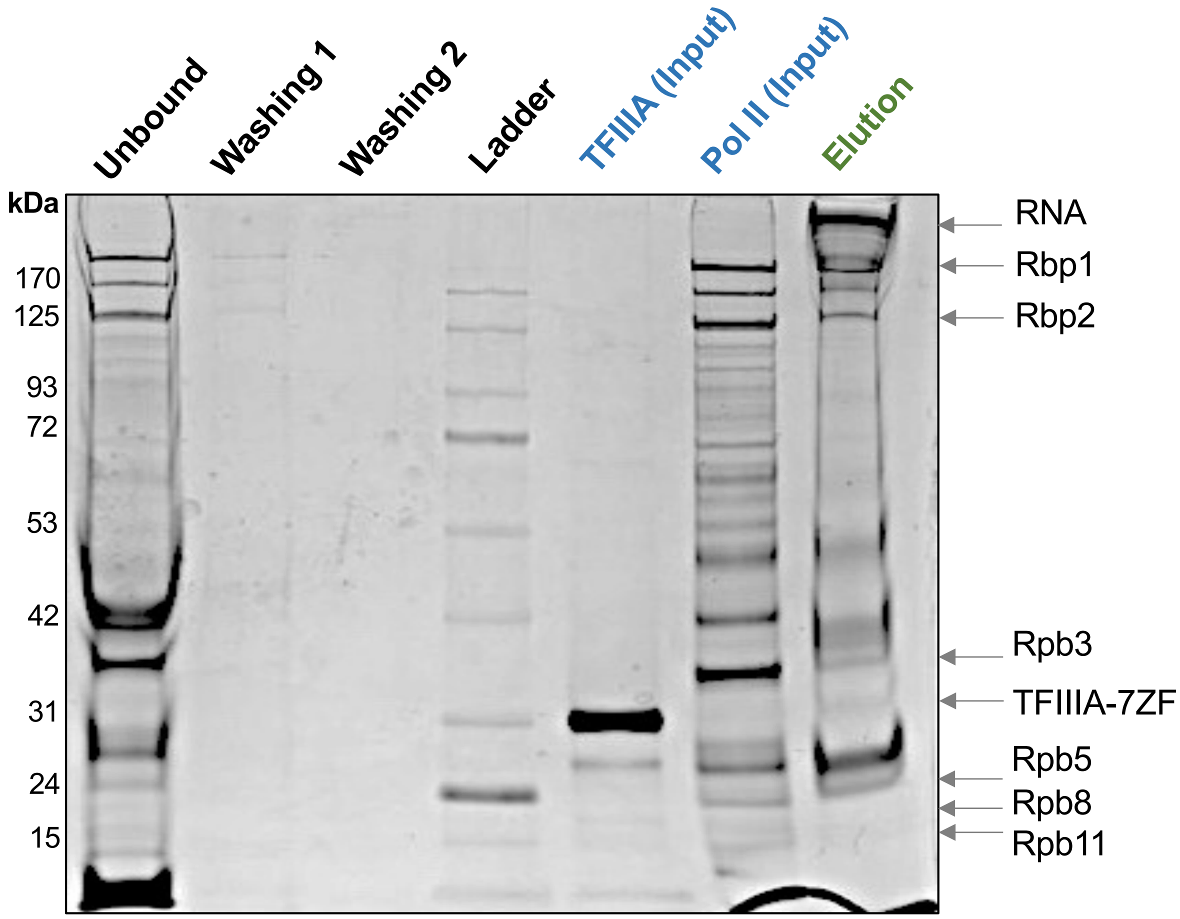


**S2 Figure.** Silver staining of RNA-based affinity purification fractions. Pol II subunits are labeled based on the predicted molecular weight.
